# Real brains in virtual worlds: Validating a novel oddball paradigm in virtual reality

**DOI:** 10.1101/749192

**Authors:** Jonathan W. P. Kuziek, Abdel R. Tayem, Jennifer I. Burrell, Eden X. Redman, Jeff Murray, Jenna Reinen, Aldis Sipolins, Kyle E. Mathewson

**Author notes:** **Corresponding Author:** Kyle E. Mathewson, Ph.D., Assistant Professor of Cognitive Neuroscience, Department of Psychology, P-455 Biological Sciences, University of Alberta, Edmonton, AB, Canada, T6G 2E9.

## Abstract

Electroencephalography (EEG) research is typically conducted in controlled laboratory settings. This limits the generalizability to real-world situations. Virtual reality (VR) sits as a transitional tool that provides tight experimental control with more realistic stimuli. To test the validity of using VR for event-related potential (ERP) research we used a well-established paradigm, the oddball task. For our first study, we compared VR to traditional, monitor-based stimulus presentation using visual and auditory oddball tasks while EEG data was recorded. We were able to measure ERP waveforms typically associated with such oddball tasks, namely the P3 and earlier N2 components, in both conditions. Our results suggest that ERPs collected using VR head mounted displays and typical monitors were comparable on measures of latency, amplitude, and spectral composition. In a second study, we implemented a novel depth-based oddball task and we were able to measure the typical oddball-related ERPs elicited by the presentation of near and far stimuli. Interestingly, we observed significant differences in early ERPs components between near and far stimuli, even after controlling for the effects of the oddball task. Current results suggest that VR can serve as a valid means of stimulus presentation in novel or otherwise inaccessible environments for EEG experimentation. We demonstrated the capability of a depth-based oddball in reliably eliciting a P3 waveform. We also found an interaction between the depth at which objects are presented and early ERP responses. Further research is warranted to better explain this influence of depth on the EEG and ERP activity.

## 1. Introduction

Electroencephalography (EEG) research is typically conducted in a highly controlled laboratory setting. However, this often limits the generalizability of results to real-world situations. Contemporary research has shown that alternative means of stimulus presentation such as virtual or augmented reality can yield results comparable to traditional EEG methods. In order to identify ways to make EEG experimentation more accessible and generalizable, here we test the use of a virtual reality head mounted display to elicit ERP components of interest in an oddball task.

Recently there has been an increase in EEG experimentation moving into more ecologically relevant environments. With this increase, there have been several notable attempts to find alternative means of stimulus presentation. Recent research into mobile EEG (Kuziek, Shienh, & Mathewson (2017); Scanlon, Townsend, Cormier, Kuziek, & Mathewson (2019); Debener, Minow, Emkes, Grandras, & De Vos (2012); has demonstrated methods to increase experimental generalizability by bringing EEG techniques into new environments. While mobile EEG experimentation is valuable and addresses many challenges facing modern cognitive neuroscience, it can be limited by such practical boundaries as participant safety in potentially hazardous environments and by the presentation of certain types of stimuli remaining unfeasible. Virtual Reality (VR) allows for the presentation of otherwise inaccessible environments while retaining a high level of experimental flexibility and control.

VR has over two decades of history in research, having been incorporated into psychology, neuroscience, and medical experimentation (Pugnetti et al., 1998; Sawaragi & Horiguchi, 1999; Lee et al., 2003). However, with recent advances in VR technology, techniques utilizing VR research are rapidly changing. VR has recently seen use in a wide range of research extending from remediation following stroke (Choi, Ku, Lim, Kim, & Paik, 2016; Ho et al., 2019), to treatment for fear of public speaking (Safir, Wallach, & Bar-Zvi, 2012; Anderson et al., 2013), and PTSD (Rothbaum et al., 2014; Smith et al., 2015). Other studies have begun to employ the use of VR in cognitive neuroscience, allowing for the creation of new paradigms, most notably in embodied cognition to which there are few other methods available to elicit relevant participant states (Argelaguet, Hoyet, Trico, & Lécuyer, 2016). Recently, VR has been increasingly used with concurrent brain recording, studies into the effects of VR have primarily used EEG as its measurement of effects. Recent studies such as Banaei et al., (2017) and Stolz, Endres, and Mueller (2019), have investigated VR while using mobile EEG to create safer immersive environments to investigate and explore.

VR research is still in its infancy; as such, there are few paradigms in research that have been appropriately validated. Dankert, Heil and Pfeiffer (2013) studied stereo vision and acuity in VR. In the study, they used VR to present animated dots to the participants, using a concave screen and projector to create a sense of immersion, and those with stereo vision were able to see a shape emerging. However, the limitations of the screen size allowed for pixilation to occur, limiting the validity of the results. The impact of VR hardware itself on a given experiment remains unclear. Results such as those from Dankert, Heil, & Pfeiffer (2013), as well as others, demonstrate the lack of systematic testing of the combination of immersive VR, projective VR, and EEG. Harjunen, Ahmed, Jacucci, Ravaja and Spapé (2017) attempted to investigate the differences between VR and non-VR conditions, concluding that “VR headsets can safely be used in ERP research without compromising the reliability of EEG recordings.” However, their conclusion was based on the lack of “adverse effects” to the ERPs, not the unexpected seemingly auspicious results of an increased P3 and signal-to-noise-ratio. Other studies such as Strickland and Chartier (1997) and Bayliss and Ballard (2000) investigated the effects of VR on EEG data. However, due to the great leaps in technology, as well as the decrease in size and weight of VR systems over the past 20 years, these papers are no longer representative and testing with modern equipment is needed for further validation.

Our goal of our first experiment (Exp. 1) was to establish the efficacy of using current VR technology and EEG simultaneously, to further validate this methodology. We did this by having participants engage in identical auditory and visual oddball tasks while wearing a VR headset or while using traditional non-VR methods. To further explore the uses of VR, we conducted a second experiment (Exp. 2) in which our goal was to demonstrate that a novel VR-specific depth-based oddball task can elicit the typical oddball-related P3 response. We were also interested to see whether there are discernible differences in ERPs to stimuli shown at near versus far depths.

## 2.0 Method - Exp. 1

### 2.1 Participants

A total of 24 members of the university community participated in the experiment (mean age = 23.33; age range =18-32; 9 males). Participants were all right-handed, and all had normal or corrected normal vision and no history of neurological problems. All participants gave informed consent and were either given course credit for an introductory psychology course or else were given an honorarium of $10/hour for their time. The experimental procedures were approved by the internal Research Ethics Board of the University of Alberta.

### 2.2 Materials & Procedure

A virtual environment was displayed to the participant with either a ViewPixx monitor (non-VR condition) or an HTC VIVE virtual reality headset (VR condition). Participants completed four oddball tasks, an auditory and a visual oddball task in each of the VR and non-VR environments. Task order, auditory or visual oddball, was counterbalanced and randomised. The order of the stimulus presentation, non-VR or VR, was counterbalanced and alternated between participants such that participant 1 would complete non-VR first, participant 2 would complete VR first, participant 3 would complete non-VR first, and so on. Participants would complete both VR or non-VR tasks before switching to the second condition (eg. both the auditory and visual oddball tasks would be completed in VR followed by completing both oddball tasks in non-VR). During each trial either a target or standard stimulus was presented, with a 20% chance for target to be presented and an 80% chance for a standard to be presented. Participants were asked to sit still and fixate on a white cross presented directly in front of them that stayed constant throughout the auditory task. In the auditory task two tones at different frequencies were played, 1000 Hz for standard tones and 1500 Hz for target tones. Each tone was sampled at 44100 Hz, presented for a duration of 16 ms and contained a 2 ms linear ramp up and down. For the visual oddball task participants were again told to focus on the fixation cross while also attending to two black orbs located to the left and right of fixation. At the beginning of each trial, these orbs would flash one of two colours, either blue for targets or green for standards. The orbs would remain coloured for approximately 1.4 s until returning to black. Participants were instructed to move only their right hand to press the spacebar on a keyboard placed in front of them each time a target was presented. Following the presentation of a standard, participants were instructed to withhold any response. For both the auditory and visual oddball tasks, the time between each stimulus onset was randomly selected from an evenly distributed array of times between 1.5-2 s. Response times were collected until 1.4 s following target onset. Figure 1A and 1B show how the auditory and visual oddball tasks were presented, respectively.

**Figure 1.**
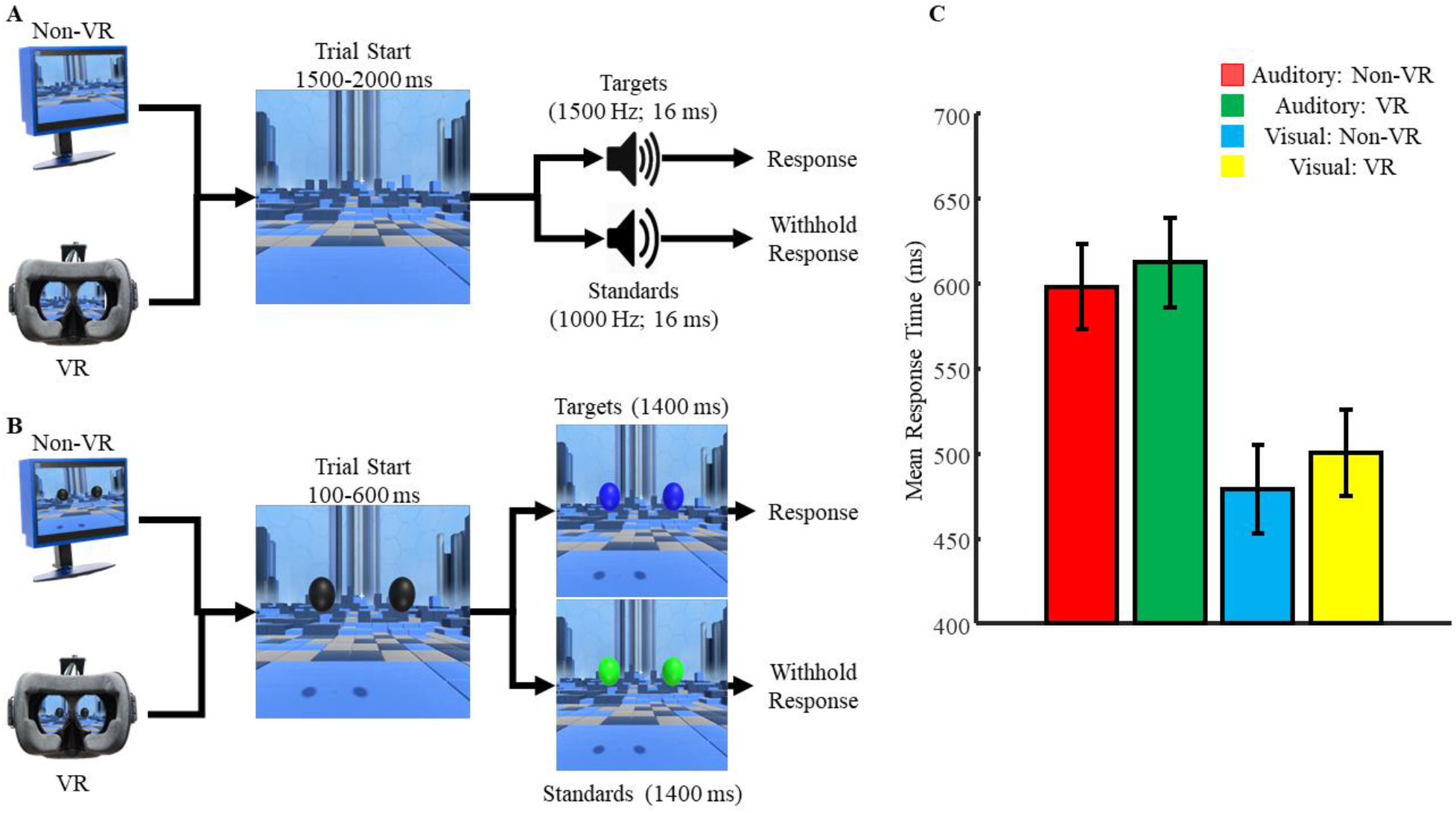
Design of the experiments used for the A) auditory oddball and the B) visual oddball tasks. C) Response times (ms) following targets during the auditory and visual oddball tasks.

For the VR condition, the size of the fixation cross was (0.02 m, 0.046 m, 0.02 m) and location was (−0.2 m,1.45 m, −0.275 m), reported in (x, y, z) coordinates with the default units in Unity being meters. The program and source code can be accessed freely from the following repository: github.com/APPLabUofA/VR_Depth. The size of each orb was (0.25 m, 0.25 m, 0.25m). The location of the left orb was (−0.58 m, 1.45 m, −0.275 m) and the location of the right orb was (0.18 m, 1.45 m, −0.275 m). The location of the participant in the virtual environment was approximately (−0.189 m, 1.3302 m, − 1.5664 m). The size of the fixation cross was approximately (0.02 m, 0.04 m, 0.02 m), located in the center of our environment (0 m, 0 m, 0 m). For the non-VR condition, the fixation measured 1.1 by 1.1 degrees and was presented in the center of the screen. Each orb was 5.9 degrees tall by 5.6 degrees wide, presented 8.5 degrees to the left and right of fixation. Stimuli during the VR condition were displayed using an HTC VIVE headset which contains two PenTile OLED displays with a maximum resolution of 2160 x 1200 pixels and a refresh rate of 90 Hz. For the non-VR condition participants were seated 57-cm away from a ViewPixx/EEG LED monitor running at a resolution of 1920 x 1080 pixels and a refresh rate of 120 Hz with simulated-backlight rastering. Each participant completed 250 trials for each oddball experiment and condition. Stimuli were presented using a Windows 7 PC running Unity 2017.1.0. Auditory stimuli were presented using a pair of Logitech Z130 speakers kept at a constant volume level. Video output was via an Asus Striker GTX1070, and audio was output via an Asus Xonar DSX sound card. Coincident in time with sound and image onset, 8-bit TTL pulses were sent to the EEG amplifier by a parallel port cable connected to the ViewPixx monitor to mark the data for ERP averaging. The experiment was displayed to the ViewPixx monitor in both conditions to allow the experimenter to confirm the task had properly started at the beginning of each condition, and to send TTL pulses to mark stimulus onset within the recorded EEG data. We tested the delay between stimulus onset and TTL pulse onset for the auditory and visual oddball tasks. While there was a noticeable delay for the auditory task (Mean = 169.5 ms; SD = 16.1 ms), compared to the visual task (Mean_VR_ = 4.8 ms; SD_VR_ = 3.0 ms; Mean_Non-VR_ = 12.9 ms; SD_Non-VR_ = 1.4 ms), these delays are consistent. We believe the auditory delays are due to auditory jitter introduced by the Windows and Unity software. Please note that auditory delays were not measured separately for the VR and non-VR conditions because the auditory stimuli were presented in Unity for both conditions.

### 2.3 EEG recording

During each oddball task, EEG data was collected from participants using passive, wet, low-impedance electrodes (actiCAP passive electrodes kept below 5 kΩ). Inter-electrode impedances were measured at the start of each experiment. All electrodes were arranged in the same 10-20 positions (O1, P7, T7, P3, C3, F3, Pz, Cz, Fz, P4, C4, F4, P8, T8, O2). A ground electrode was used, positioned at AFz. Ag/AgCl disk electrodes were used, with SuperVisc electrolyte gel and mild abrasion with a blunted syringe tip used to lower impedances. Gel was applied and inter-electrode impedances were lowered to less than 5 kΩ for all electrode sites. Electrode impedance was checked several times throughout the experiment; following placement of the electrodes and EEG cap on the participant, after the VR headset was placed on the participant, and after each oddball task was completed. EEG was recorded online and referenced to an electrode attached to the left mastoid. Offline, the data were re-referenced to the arithmetically derived average of the left and right mastoid electrodes.

EEG data was recorded with a Brain-Amp 16-channel amplifier connected to a laptop running the BrainVision Recorder software, using identical settings across all participants. In addition to the 15 EEG sensors, two reference electrodes, and the ground electrode, vertical and horizontal bipolar EOG was recorded from passive Ag/AgCl easycap disks. EOG electrodes were affixed vertically above and below the left eye and affixed horizontally 1 cm lateral from the outer canthus of each eye. The participant’s skin was cleaned using Nuprep (an exfoliating cleansing gel) before the placement of the electrodes, electrolyte gel was used to lower the impedance of these electrodes to under 5 kΩ in the same manner as previously mentioned. These bipolar channels were recorded using the AUX ports of the Brain-Amp amplifier. Data were digitized at 1000 Hz with a resolution of 24 bits and hardware filtered online with a low cut-off of 10.0 s, a high cut-off of 250 Hz, an additional online software cut-off of 50Hz, and a time constant of 1.59155 s. Each experiment was completed in a dimly lit, sound and radio frequency-attenuated chamber from Electromedical Instruments, with copper mesh covering the window. The only electrical devices in the chamber were an amplifier, speakers, keyboard, mouse, monitor, and VR headset. The monitor and HTC VIVE headset ran on DC power from outside the chamber, the keyboard and mouse were plugged into USB extenders outside the chamber, and the speakers and amplifier were both powered from outside the chamber. The HTC VIVE system’s two motion sensors, known as lighthouses, are mounted in the chamber in front of and to the left of the participant, as well as behind and to the right, and are raised 3 feet above the average participant’s head level. They were connected to power outside of the chamber, and they emit infrared pulses at 60Hz which are detected by the VIVE headset to locate itself in 3D space. The fan for the chamber was turned on, and nothing was plugged into the internal power outlets. Any devices transmitting or receiving radio waves (e.g., cell phones) were either turned off or removed from the chamber for the duration of the experiment.

### 2.4 EEG analysis techniques

Analyses were computed in Matlab R2018a using EEGLAB (Delorme & Makeig, 2004) and custom scripts. The timing of the TTL pulse was marked in the recorded EEG data and used to construct 1200 ms epochs time locked to the onset of standard and target tones with the average voltage in the first 200 ms baseline period, subtracted from the data for each electrode and trial. To remove artifacts due to amplifier blocking and other non-physiological factors, any trials with a voltage difference from baseline larger than +/− 1000 μV on any channel (including eyes) were removed from further analysis. At this time, a regression-based eye-movement correction procedure was used to estimate and remove the artifact-based variance in the EEG due to blinks as well as horizontal and vertical eye movements (Gratton, Coles, & Donchin, 1983). After identifying blinks with a template-based approach, this technique computes propagation factors as regression coefficients predicting the vertical and horizontal eye channel data from the signals at each electrode. The eye channel data is then subtracted from each channel, weighted by these propagation factors, removing any variance in the EEG predicted by eye movements. Artifact rejection was again performed except for this second round of artifact rejection removed trials containing a voltage difference of +/− 500μV. Baseline correction was performed a second time following the second artifact rejection.

#### 2.4.1 ERP analysis

Trial-averaged ERPs were derived and waveforms were compared. Analysis on the N2 and P3 waveforms was done using difference waveforms, computed by subtracting the ERPs for standards from targets at electrodes Fz and Pz, respectively. Appropriate time window cut-offs for the N2 and P3 waveforms were determined by running a one-tailed t-test across each time point of the participant grandaverage ERP, with α set to 0.005, to determine large, consistent regions of significance. This was only done using the ERPs derived from the non-VR condition as we used this condition as the standard of comparison for the VR-derived ERPs. Based on this technique we used the following time windows for all further analysis: Auditory N2 = 324-366 ms; Auditory P3 = 427-631 ms; Visual N2 = 88-106 ms; Visual P3 = 300-570 ms. A one-tailed t-test was performed to understand if a significant N2 or P3 peak was observed, while a two-tailed t-test was performed to compare peaks across devices, with α set to 0.05 for all analyses. We also employed Bayesian statistics, which has been proposed as a more-informative alternative to statistical tests involving p-values (Jarosz & Wiley, 2014; Wagenmakers, Love, et al., 2018; Wagenmakers, Marsman, et al., 2018) to directly test our null hypothesis that there is no significant difference in the EEG recordings between our devices. Bayesian analysis was done using version 0.8.6 of the JASP software (JASP Team, 2019), which is a powerful tool for Bayesian statistics (Wagenmakers, Love, et al., 2018; Wagenmakers, Marsman, et al., 2018).

#### 2.4.2 Single trial noise

We then estimated the amount of noise in the data on individual trials in two ways. First, we computed the average frequency spectra of the baseline period in each EEG epoch. For each participant, we randomly selected 180 of their artifact-free standard trials from electrode Pz. For each trial, we computed a Fast Fourier Transform (FFT) by symmetrically padding the 1200 time-point epochs with zero to make a 2048-point time series for each epoch, providing frequency bins with a resolution of 0.488 Hz. Each participant’s 180 spectra are then averaged together to compute participant spectra, which were then combined to form grand average spectra. To compute a second and related estimate of the noise on single trial EEG epochs, we randomly selected 180 standard tone epochs for each participant and computed the root mean square (RMS) of the baseline period on each trial. We used the 200 ms baseline period (200 time points) prior to trigger onset to avoid the influence of any evoked ERP activity on the RMS measurement. The RMS is a measure of the average absolute difference of the voltage around the baseline and is a good estimate of single trial noise in the EEG data. Smaller RMS values would suggest fewer voltage differences around the baseline and, therefore, less noise present in the data. For each trial, we average the RMS values for each EEG electrode, then averaged over trials for each participant, then computed the grand average RMS across participants (as in Laszlo, Ruiz-Blondet, Khalifian, Chu, & Jin, 2014). To estimate the distribution of RMS in our data for each device, we employed a permutation test in which 180 different epochs were selected without replacement for each participant on each of 10,000 permutations. For each of these random selections, and each electrode, we computed and recorded the grand average single trial RMS. To quantify the level of noise in the participant average ERPs, we again employed a permutation test of the RMS values in the baseline period. In this ERP version, for each of the 10,000 permutations, we averaged the 180 standard trials that were randomly selected without replacement from the larger pool of that participant’s artifact free trials for each device. We then computed the RMS of the resultant 200 time points of the ERP baseline. We averaged these RMS values over all EEG electrodes, then computed a grand average across participants.

#### 2.4.3 ERP power

To compare the ERP statistical power as a function of the number of trials used for both the P3 and N2, we used another permutation procedure in which we varied the number of trials contributing to the ERP average while keeping the 4 to 1 ratio of standard to target trials (Mathewson, Harrison, & Kizuk, 2017). Trial numbers were varied from 4 standards and 1 target trial, by 20 standard trials, up to 180 standard and 36 target trials, separately for each of the two stimulus types. For each number of trials, 10,000 permutations were randomly selected from the total pool without replacement. For each permutation, the selected single trials were averaged to create participant ERPs separately for target and standard tones. The difference between target and standard tones was then computed at electrode Fz (N2) and electrode Pz (P3), and these simulated participant average ERP differences were compared to a null distribution with a standard one-tailed t-test.

## 3.0 Results - Exp. 1

### 3.1 Behavioural Results

Following artifact rejection similar trial counts were obtained for targets and standards as shown in Table 1. Figure 1C shows the response times to target stimuli during the auditory and visual oddball tasks. Results of a two-tailed t-test suggest no difference in response times between the auditory VR (Mean = 612.521 ms; SD = 128.375 ms) and non-VR conditions (Mean = 598.212 ms; SD = 123.112 ms; t(23) = −0.519; p = 0.609) but a significant difference between the visual VR (Mean = 500.293 ms; SD = 124.616 ms) and non-VR conditions (Mean = 479.360 ms; SD = 127.372 ms; t(23) = −2.476; p = 0.021).

**Table 1:**
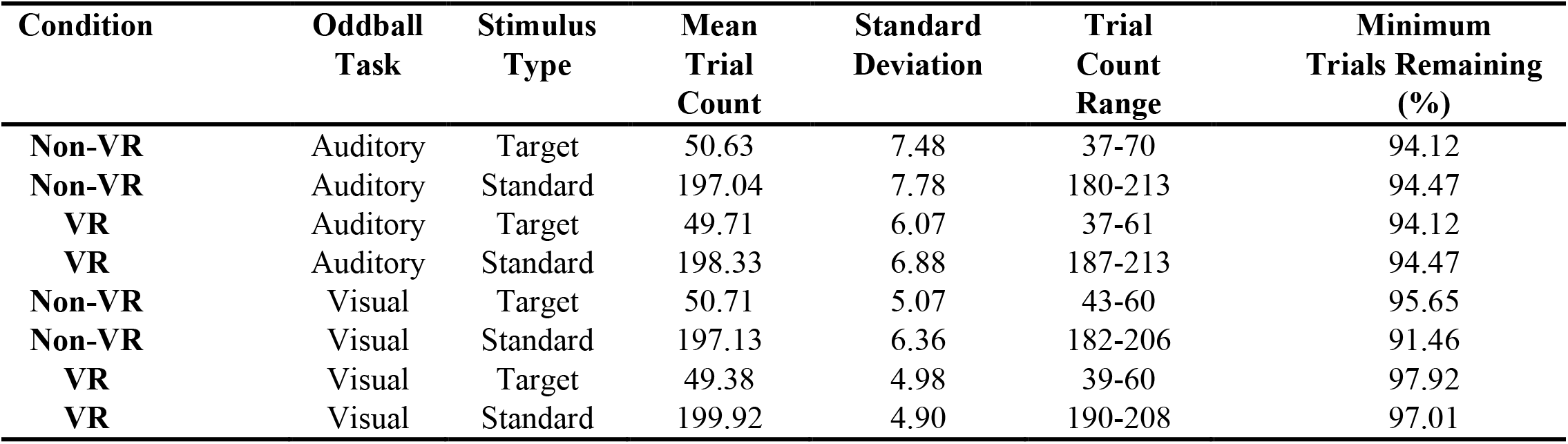
Trial count information for both target and standard trials across condition (Non-VR; VR) and oddball modality (Auditory; Visual).

| Condition | Oddball Task | Stimulus Type | Mean Trial Count | Standard Deviation | Trial Count Range | Minimum Trials Remaining (%) |
| --- | --- | --- | --- | --- | --- | --- |
| Non-VR | Auditory | Target | 50.63 | 7.48 | 37-70 | 94.12 |
| Non-VR | Auditory | Standard | 197.04 | 7.78 | 180-213 | 94.47 |
| VR | Auditory | Target | 49.71 | 6.07 | 37-61 | 94.12 |
| VR | Auditory | Standard | 198.33 | 6.88 | 187-213 | 94.47 |
| Non-VR | Visual | Target | 50.71 | 5.07 | 43-60 | 95.65 |
| Non-VR | Visual | Standard | 197.13 | 6.36 | 182-206 | 91.46 |
| VR | Visual | Target | 49.38 | 4.98 | 39-60 | 97.92 |
| VR | Visual | Standard | 199.92 | 4.90 | 190-208 | 97.01 |

### 3.2 ERP Analysis

Figure 2 shows the grand average ERPs from electrode Pz and Fz following standard and target tones during the A) auditory and B) visual oddball tasks. A P3 response can be observed following targets in both oddball tasks but an N2 response is only strongly observed following auditory targets. Also shown are the difference waveforms at electrode Fz and Pz for the A) auditory and B) visual oddball tasks. The difference waveforms were calculated by subtracting the ERPs for standards from the ERPs following targets. A large negative deflection at electrode Fz can be observed for the auditory N2. By comparison, a much smaller negative deflection is observed for the visual N2. A distinct P3 response, signified by a large positive-going deflection, can be observed at electrode Pz for both the auditory and visual oddball tasks. Figure 2 also shows topographies and bar graphs of the averaged ERP effects across the indicated time windows, with similar patterns of activity for both devices in all but the visual N2 region.

**Figure 2:**
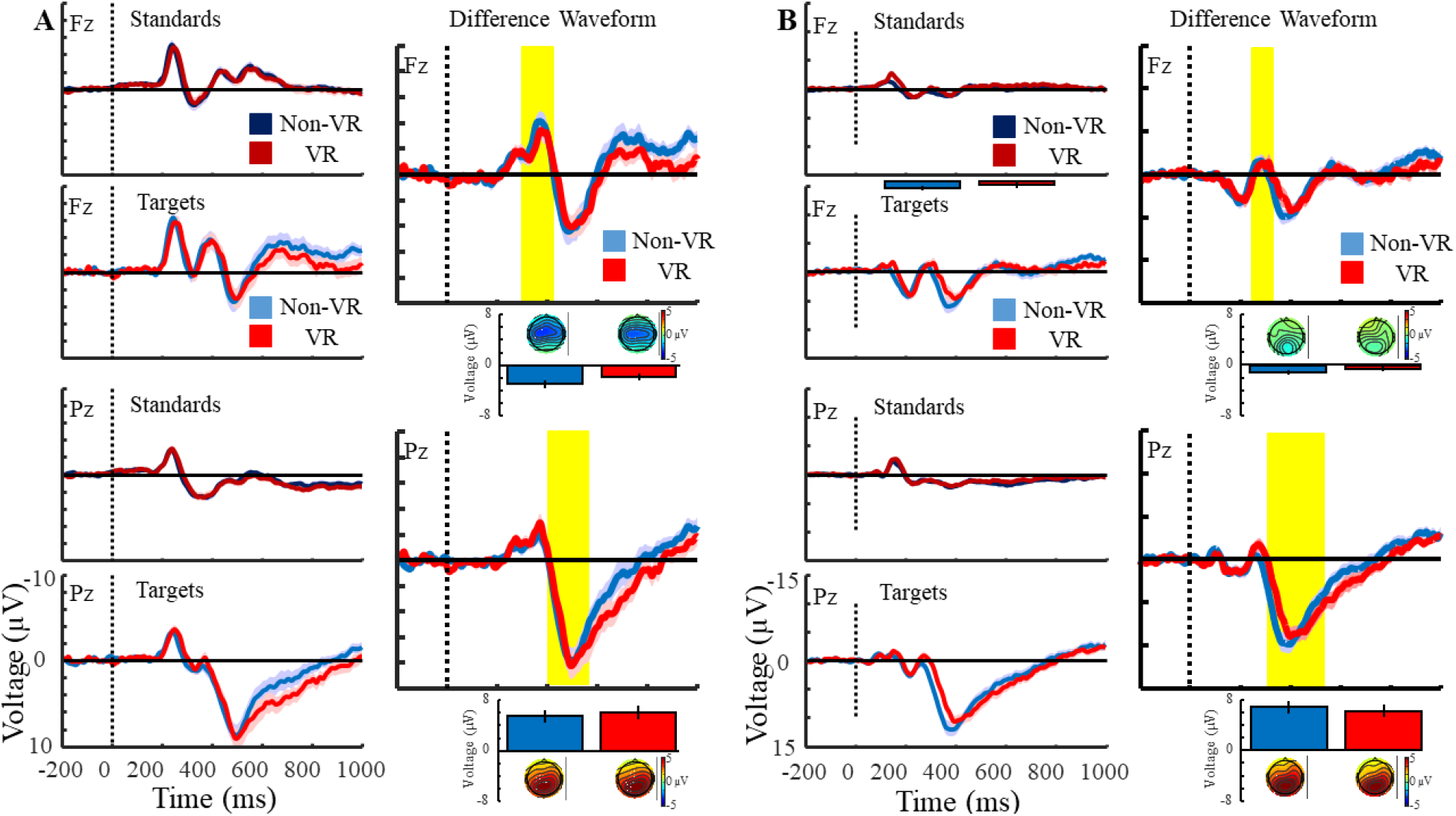
Grand-averaged ERP plots for target and standard stimuli at electrode Fz and Pz. Also shown are the difference waveforms for the A) auditory oddball and B) visual oddball tasks at electrode Fz and Pz. Difference waveforms were generated by subtracting the ERPs following standard stimuli from the ERPs following target stimuli. Yellow shaded regions represent the N2 and P3 time windows used for analyses. Topographies represent activity across the scalp while the bar graphs show activity at either electrode Fz or Pz, averaged across the N2 and P3 time windows. Error bars represent the standard error of the mean.

Table 2A shows the results of the one-tailed *t*-test for the N2 and P3 waveforms at electrode Fz and Pz respectively. Results comparing the waveforms between the VR and non-VR condition are shown in Table 2B. Hedge’s *g* was used to estimate effect size, which was calculated using version 1.5 of the Measures-of-Effect-Size toolbox for Matlab (Hentschke & Stüttgen, 2011). Results of the Bayesian one-sample and paired-sample t-tests show how much our data supports the alternative hypothesis for each test, either being a significant difference from 0 for the one-sample t-tests of a significant difference between the VR and non-VR conditions for the paired-sample t-tests. Support for the alternative hypothesis is reported as a Bayes Factor (BF10) with numbers greater than 1 indicating the data is more likely under the alternative hypothesis and numbers less than 1 indicating support for the null hypothesis.

**Table 2:** Results of one-tailed and two-tailed *t*-tests, Bayesian one-tailed and two-tailed *t*-tests, and estimated measures of effect size on the ERP difference waves. A) One tailed *t*-test and Bayes Factor results to determine if a significant N2 and P3 response was observed for each condition and oddball task. Bayes factors were calculated by testing if the N2/P3 waveforms for each condition/oddball task were less than or greater than zero, respectively. Hedge’s *g* is used to estimate effect size. B) Two-tailed *t*-test and Bayes paired-t-test results comparing the N2 and P3 waveforms for each condition and oddball task. Bayes Factors (BF10) were calculated by comparing the N2 and P3 waveforms between Non-VR and VR conditions.

| Condition | Oddball Task | Electrode/<br>Time Window | Mean Voltage<br>(μV) | Standard Deviation<br>(μV) | $t$ (23) | $p$ | Confidence Interval | Hedge's $g$ | $BF_{10}$ |
| --- | --- | --- | --- | --- | --- | --- | --- | --- | --- |
| Non-VR | Auditory | Fz/N2 | -2.98 | 2.71 | -5.40 | <0.01 | -Inf:-2.03 | -1.07 | 1.30e3 |
| Non-VR | Auditory | Pz/P3 | 5.50 | 4.20 | 6.42 | <0.01 | 4.04:Inf | 1.27 | 1.24e4 |
| VR | Auditory | Fz/N2 | -1.88 | 2.77 | -3.32 | <0.01 | -Inf:-0.91 | -0.66 | 13.52 |
| VR | Auditory | Pz/P3 | 6.06 | 5.08 | 5.84 | <0.01 | 4.28:Inf | 1.15 | 3.51e3 |
| Non-VR | Visual | Fz/N2 | -0.03 | 1.53 | -0.10 | 0.46 | -Inf:0.50 | -0.02 | 0.22 |
| Non-VR | Visual | Pz/P3 | 6.58 | 3.77 | 8.55 | <0.01 | 5.26:Inf | 1.69 | 9.72e5 |
| VR | Visual | Fz/N2 | 0.33 | 1.94 | 0.83 | 0.79 | -Inf:1.01 | 0.16 | 0.30 |
| VR | Visual | Pz/P3 | 6.00 | 3.72 | 7.89 | <0.01 | 4.70:Inf | 1.56 | 2.65e5 |

| Oddball Task | Electrode/<br>Time Window | Mean Voltage<br>(μV) | Standard Deviation<br>(μV) | $t$ (23) | $p$ | Confidence Interval | Hedge's $g$ | $BF_{10}$ |
| --- | --- | --- | --- | --- | --- | --- | --- | --- |
| Auditory | Fz/N2 | -1.10 | 3.35 | -1.40 | 0.17 | -2.70:0.49 | -0.39 | 0.67 |
| Auditory | Pz/P3 | -0.55 | 2.50 | -0.41 | 0.68 | -3.26:2.16 | -0.12 | 0.36 |
| Visual | Fz/N2 | -0.36 | 2.33 | -0.71 | 0.48 | -1.37:-0.71 | -0.20 | 0.28 |
| Visual | Pz/P3 | 0.59 | 2.66 | 0.54 | 0.59 | -1.59:2.77 | 0.15 | 0.36 |

### 3.3 Single Trial Noise

Grand average spectra are shown in Figure 3A and 3C, along with topographies showing the average spectra between 0.5-3 Hz, 4-7 Hz, 8-12 Hz, and 13-30 Hz. Histograms of the single trial RMS values computed for each permutation, along with bar graphs of the mean and standard deviation, are shown in Figure 3B and 3E for the auditory and visual oddball tasks respectively. Also shown are the grand average RMS values computed in each of the 10,000 permutations for both conditions, along with a bar graph of the mean and standard deviation. Both the single trial and grand average RMS distributions suggest less noise was present during the non-VR condition compared to the VR condition.

**Figure 3:**
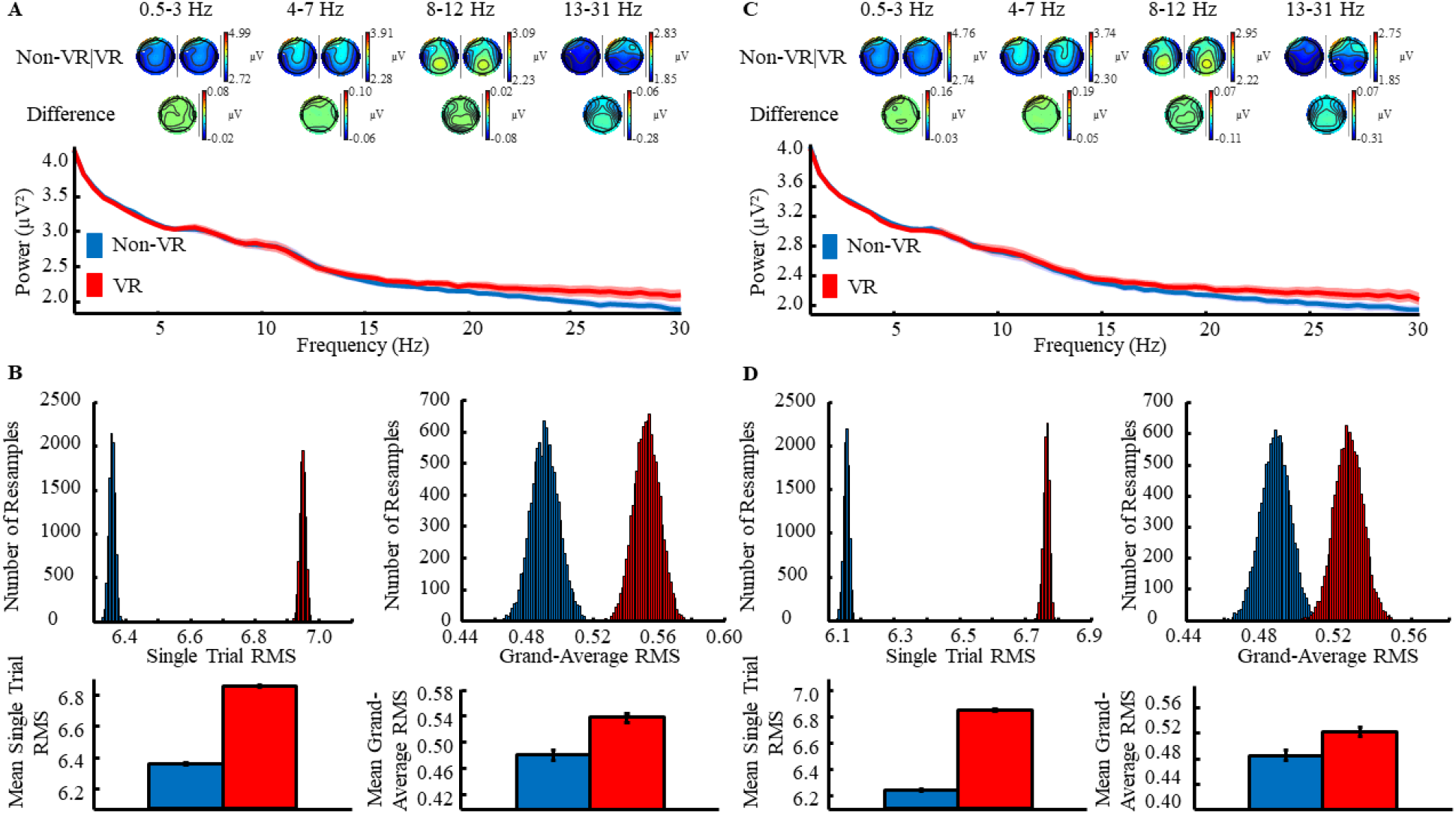
Spectra and RMS analysis for the auditory and visual oddball tasks. A) Grand-averaged spectra, topographies, and B) single-trial and grand-averaged RMS distributions for the auditory oddball task. Error bars in the spectra plots represent the standard deviation at each frequency. C) Grand-averaged spectra, topographies, and D) single-trial and grand-averaged RMS distributions for the oddball task. Error bars on the mean RMS plots represent the standard deviation of the respective RMS distribution.

### 3.4 ERP Power

Figure 4 plots the proportion of the 10,000 permutations in which the *t*-statistic passed the significance threshold, as a function of the number of samples in each permutation. The P3 and N2 waveforms required a similar number of trials to reach significance with 80% power, although as expected the N2 waveform did not reach significance for either condition during the visual oddball task. For the N2 waveform during the auditory task, approximately 37 and 26 trials were needed for the VR and non-VR conditions respectively. For the P3 waveform during the auditory task about 6 trials were needed to reach significance for both conditions. For the visual oddball task about 4 trials were needed for the P3 waveform to reach significance in both conditions.

**Figure 4:**
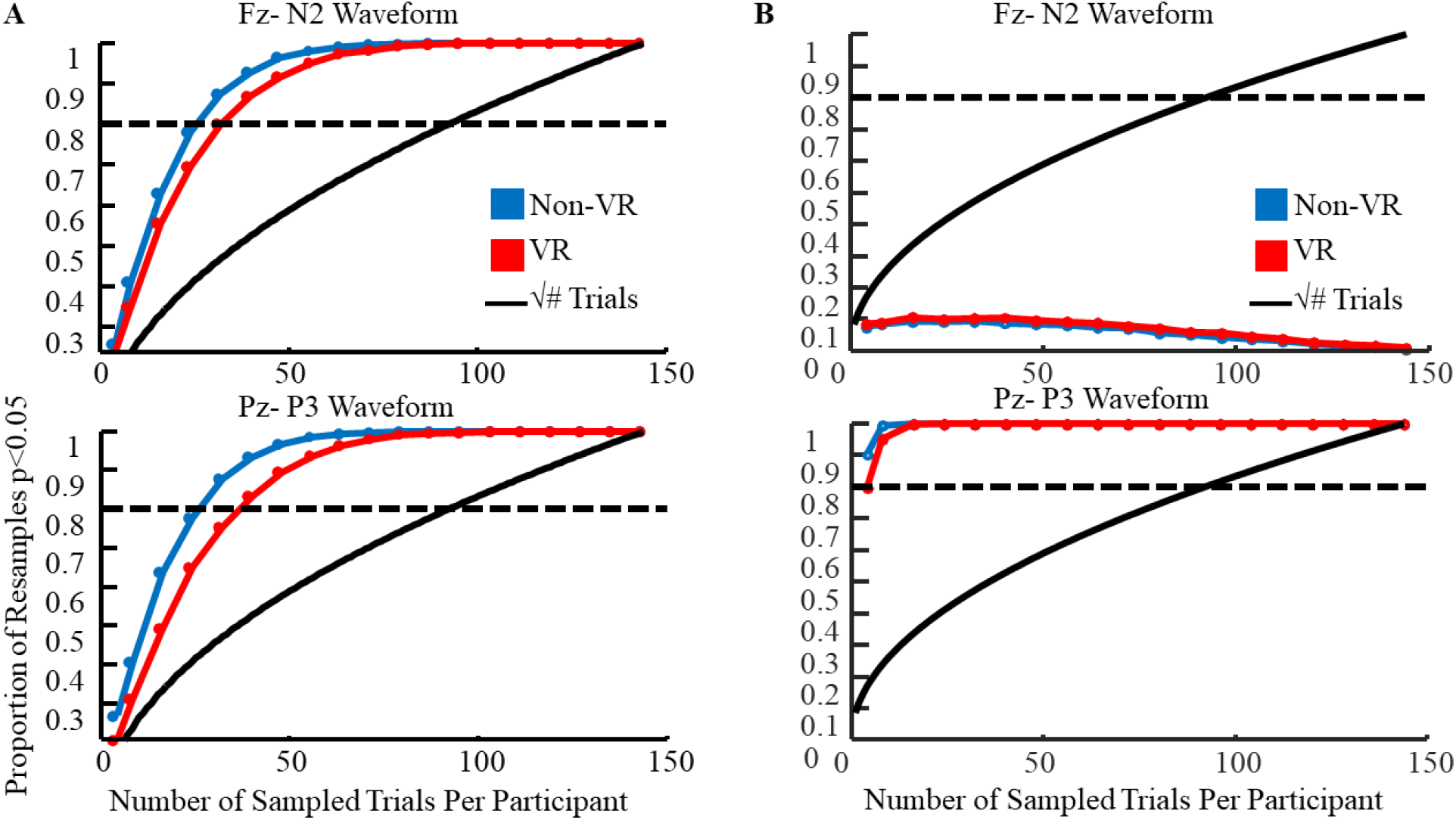
Plots showing the trials counts needed to obtain a significant N2 or P3 with 80% power for the A) auditory oddball and B) visual oddball tasks. Analyses were run using electrode Fz for the N2 waveform and electrode Pz for the P3 waveform.

## 4. Discussion - Exp. 1

The goal of the first experiment was to expand on a recent trend in current literature focusing on EEG generalization and to establish validity in the use of VR technology in basic research in cognitive neuroscience, particularly those involving the oddball paradigm.

Behavioural results indicate there is a small but significant difference between the non-VR and VR in the visual oddball task. Overall slower responses to 3D stimuli may be the result of increased visual information, such as depth, being presented to participants. Another explanation could be the distraction of wearing a VR headset causes a slowing of response times as well. However, despite these differences, ERP waveforms derived from stimuli presented in the VR and non-VR conditions were found to be nearly identical during the time windows relevant to the oddball task. Taken together these results demonstrate the versatility of VR and its applicability in cognitive psychology. It is important to note the clarity in the P3 response despite introduced sources of noise in the VR condition. The first of which is the addition of VR equipment which may introduce electrical noise and presents an added physical challenge to participants with added weight, which may lead to extra muscle artifacts due to neck tension, or increased pressure on electrodes directly beneath the headset. The second source of noise pertains to the different levels of engagement within the environment, differing levels of processing may be required to interpret a more immersive and realistic scene, however, this may be counteracted by virtue of the VR environment itself presenting as more engaging to the viewer and yielding more readily available attentional resources. Lastly, there are several differences between non-VR and VR stimuli in terms of viewing quality: resolution, refresh rate, and field of view.

Both systems showed the expected ERP, P3, and N2 waveforms along with similar topographies, with the greatest activity focused on electrodes Pz and Fz respectively. Bayesian paired t-tests further support that it is likely the data falls under the null hypothesis suggesting no difference between recording techniques for the N2 and P3. However, we did observe some differences in the amount of noise during baseline, as indicated by the differences in the RMS distributions for both systems. Despite the apparent difference in noise between the VR and non-VR conditions, similar statistical power, frequency distributions, and ERP waveforms were observed between both systems. However, other results have shown that any observed difference between RMS noise has minimal impact on the derived ERPs and their statistical power.

Both devices showed the expected 1/f frequency structure in the spectral data, as well as the typical peak in the alpha frequency range between 8 and 12 Hz (Mathewson et al., 2011) along with another peak at about 7Hz. At frequencies below approximately 15Hz, both conditions show similar spectra but start to deviate past 15Hz. This difference could be due to several sources. The VR headset likely put pressure on several frontal electrodes, along with any electrodes directly under the headset straps. Increased facial and muscle strain was also likely present for the VR condition due to the weight and comfort of the headset. While only the muscle-related sources of noise would likely be present in the frequency range we are interested in (Hagemann & Naumann, 2001; Goncharova, McFarland, Vaughan, & Wolpaw, 2003), increased electrical sources of noise would still be present during the VR condition and likely influence frequencies far beyond our 30 Hz maximum. Furthermore, it is worth noting there is a consistent trend in the commercial VR industry to provide progressively lighter VR head mounted displays; there will likely be less contamination due to muscle artifacts with lighter models.

We were able to demonstrate that a commercial HTC Vive headset, a relatively inexpensive and fully immersive VR device, can present oddball stimuli comparably to a research-standard non-VR monitor. We were able to show that the ERPs derived from typical auditory and visual oddball tasks are comparable when stimuli are presented using VR and non-VR methods. We then looked to explore stimuli relevant to 3D environments. In the next experiment, we explore the addition of a new dimension, depth, to the standard visual oddball task, a task that cannot be replicated on a traditional monitor.

## 5. Introduction - Exp. 2

Results from Exp.1 indicate that the ERPs from stimuli presented from VR headsets are comparable to those presented with a traditional non-VR monitor. In the following experiment, we hoped to further demonstrate the utility of VR in an EEG experiment wherein the paradigm domain is more specific to VR itself. We translated the typical oddball paradigm into a novel VR-specific task by specifically modulating the domain of depth. Using this depth-based oddball, we compared ERPs to near and far standards and targets in an attempt to elicit a P3 waveform similar to other recent adaptations of the standard oddball task (Raz, Dan, & Zysberg, 2014). The oddball paradigm has been shown to be sensitive to changes in several modalities, ranging from sensitivity to basic visual elements of stimuli such as colour (Fonteneau & Davidoff, 2007), to higher-level processes such as emotion (Raz & Zysberg, 2014).

Depth has traditionally been studied by moving a stimulus or monitor while covering the participant’s eyes but does not allow for perceived size (retinal size) to remain constant. This effect contributes to the ease in which participants are able to identify the near or far object due to extraneous depth cues (Bohr & Read, 2013). When using VR, object size can be controlled proportionally to keep retinal size constant at varying depths with respect to the viewer. Other potential depth cues, such as shadows, may also be controlled in a VR environment. Naceri, Moscatelli, & Chellali (2015) studied depth perception when perceived size was not available as a depth cue. They found that while keeping the size of the objects constant, participants used convergence, binocular disparity and accommodation cues to allow for depth recognition. Our study followed this principle and kept the perceptual size consistent across different depths.

## 6. Method - Exp. 2

### 6.1 Participants

A total of 18 members of the university community participated in experiment two (mean age=21.60 age range=17-43; 8 males). Participants were all right-handed, and all had normal or corrected normal vision and no history of neurological problems. All participants gave informed consent, were given course credit for their time, and the experimental procedures were approved by the internal Research Ethics Board of the University of Alberta.

### 6.2 Materials & Procedure

All hardware remained the same as in the VR condition described in Exp.1. Each participant completed a visual oddball task in VR where orbs were placed either close to the participant (Near) or further away (Far). The size of the near orb was (0.10 m 0.10 m, 0.10 m) and the location was (−0.2 m, 1. 45 m, −0.162 m). The size of the far orb was (1.0 m, 1.0 m, 1.0 m) and the location was (−0.29 m, 2.55 m, 12.83 m). The location of the participant in the virtual environment was identical to experiment one (−0.189 m, 1.3302 m, −1.5664 m). The size of the fixation cross was approximately (0.02 m, 0.04 m, 0.02 m), located at (−0.5 m, −6.49 m, −0.04 m). The orbs participants responded to were considered targets and orbs that participants were not responding to were considered standards, in-line with the typical oddball paradigm. The experiment was divided into two conditions, a Near condition where participants responded when the nearest orbs were presented, and a Far condition where participants responded to the farthest orbs. The conditions henceforth are referred to by the location of the target stimuli Near and Far, respectively. Targets and standards were presented in the same 20% – 80% ratio as described in Exp. 1 for both Near and Far conditions.

Participants completed each condition twice, responding to near and far targets in separate blocks. Both orbs were centred and in line with the central fixation cross and both orbs were perceived to be identical in size. In other words, the farther orb was larger than the near orb but both orbs were perceived to be the same size by the participant, only the relative retinal disparity would be modulated. Participants placed their head in a chin rest to reduce noise caused by muscle strain as well as to remove extraneous depth cues due to head movement and position, such that the only distinguishing factor in the VR environment was relative depth. Task order, whether they were responding to near or far targets, was counterbalanced and alternated between participants such that participant 1 completed both Near condition blocks first, participant 2 completed the Far condition first, participant 3 completed the Near condition first, and so on. Participants were instructed to move only their right hand to press the spacebar on a keyboard placed in front of them each time a target stimulus was presented. Following the presentation of a standard stimulus, participants were instructed to withhold any response. During each trial, a fixation cross was presented for a variable duration randomly selected between 500-1000 ms, after which the fixation cross was removed and an orb was presented. Each orb was kept black and remained on screen for 1400 ms. EEG recording, analysis, artifact rejection, and eye-blink correction procedures were identical to those used in Exp.1. Figure 5A and 5B demonstrate the design of the depth oddball task. To estimate adduction and abduction of the eyes to stimuli of different depths in the task, the position of the horizontal EOGs were modified and placed over of the medial and lateral rectus muscles of the left eye.

## 7. Results - Exp. 2

### 7.1 Behavioural Results

Trial counts for targets and standards in each condition, following artifact rejection, are shown in Table 3. Figure 5C shows the mean response times following targets in the Near Targets and Far Targets conditions. Results from a two-tailed t-test suggests no difference between response times for far targets (Mean = 593.566 ms; SD = 175.966 ms) and near targets (Mean = 560.338 ms; SD = 161.841 ms; t(17) = −1.950; p =0.068).

**Figure 5:**
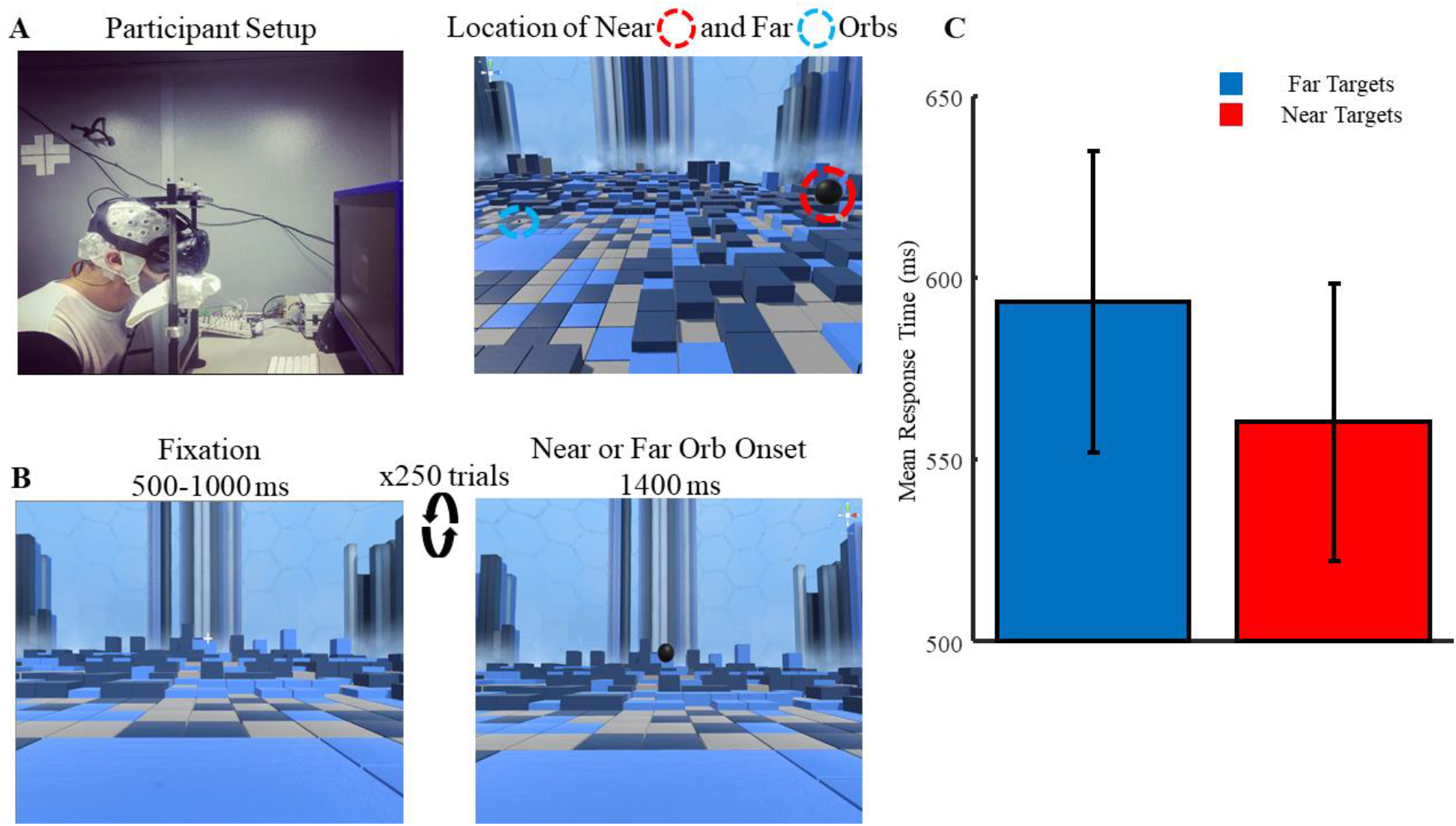
Design and setup of the depth oddball task. A) Task setup for each participant and the location of the near and far orbs with respect to the participant. B) Virtual environment and appearance of the near and far stimuli from the perspective of the participant. C) Response times for target stimuli at both near and far depths. Error bars represent the standard error of the mean.

**Table 3:** Trial count information for both target and standard trials across both the Far and Near conditions.

| Stimulus Type | Mean Trial Count | Standard Deviation | Trial Count Range (Minimum:Maximum) | Minimum Trials Remaining (%) |
| --- | --- | --- | --- | --- |
| Far Targets | 94.44 | 9.72 | 80:112 | 72.10 |
| Near Standards | 394.72 | 38.73 | 244:416 | 62.72 |
| Near Targets | 100.33 | 7.51 | 87:113 | 96.67 |
| Far Standards | 397.06 | 8.39 | 377:412 | 95.93 |

### 7.2 ERP Analysis

Figure 6A shows the grand average ERPs from electrode Pz and Fz following standard and target tones during the Near and Far conditions of the depth oddball task. A noticeable P3 response can be observed following targets in both conditions. Figure 6B shows difference waveforms for standards and targets at electrode Fz and Pz. These waveforms were constructed by subtracting Near standards from Far standards and Near targets from Far targets to observe any potential differences between Near and Far stimuli. Deflections in these difference waveforms suggest differences in ERP activity following Near/Far standards and Near/Far targets. Such deflections are typically occurring early in the difference waveform (up to 500 ms post-stimulus onset) with target stimuli showing the greatest difference in activity between the near and far depths. Figure 6C shows the difference waveforms at electrode Fz and Pz for the Near and Far conditions. These waveforms were calculated by subtracting the ERPs for standards from the ERPs following targets. In the case of the Near condition, ERPs to the far, standard stimuli were subtracted from the ERPs following the near, target stimuli. Difference waveforms for the Far condition were generated by subtracting ERPs following Near, standard stimuli from ERPs generated after the Far, target stimuli. The boundaries for the P3 time window was selected in a similar manner as described for experiment one. We ran a one-tailed t-test across each time point of the participant grand-average ERP for both the Near and Far conditions, with α set to 0.005, to determine large, consistent regions of significance. Significant regions that overlapped in both Near and Far conditions were used for our P3 time window. Based on this technique we used 349-623 ms for our P3 time window. A large P3 response during this time window at electrode Pz can be observed in the difference waveforms for both conditions. Interestingly, two distinct, opposing patterns of activity between our conditions can be seen prior to the onset of the P3 response. These patterns of activity at electrode Pz, along with the P3 waveform, were used to generate three time windows that will be used for all further analysis (175-225 ms; 225-300 ms; 349-623 ms). Electrode Pz was chosen as we expected the P3 waveform would be greatest at this electrode and, as such, any differences between our Far and Near targets would also be greatest.

**Figure 6:**
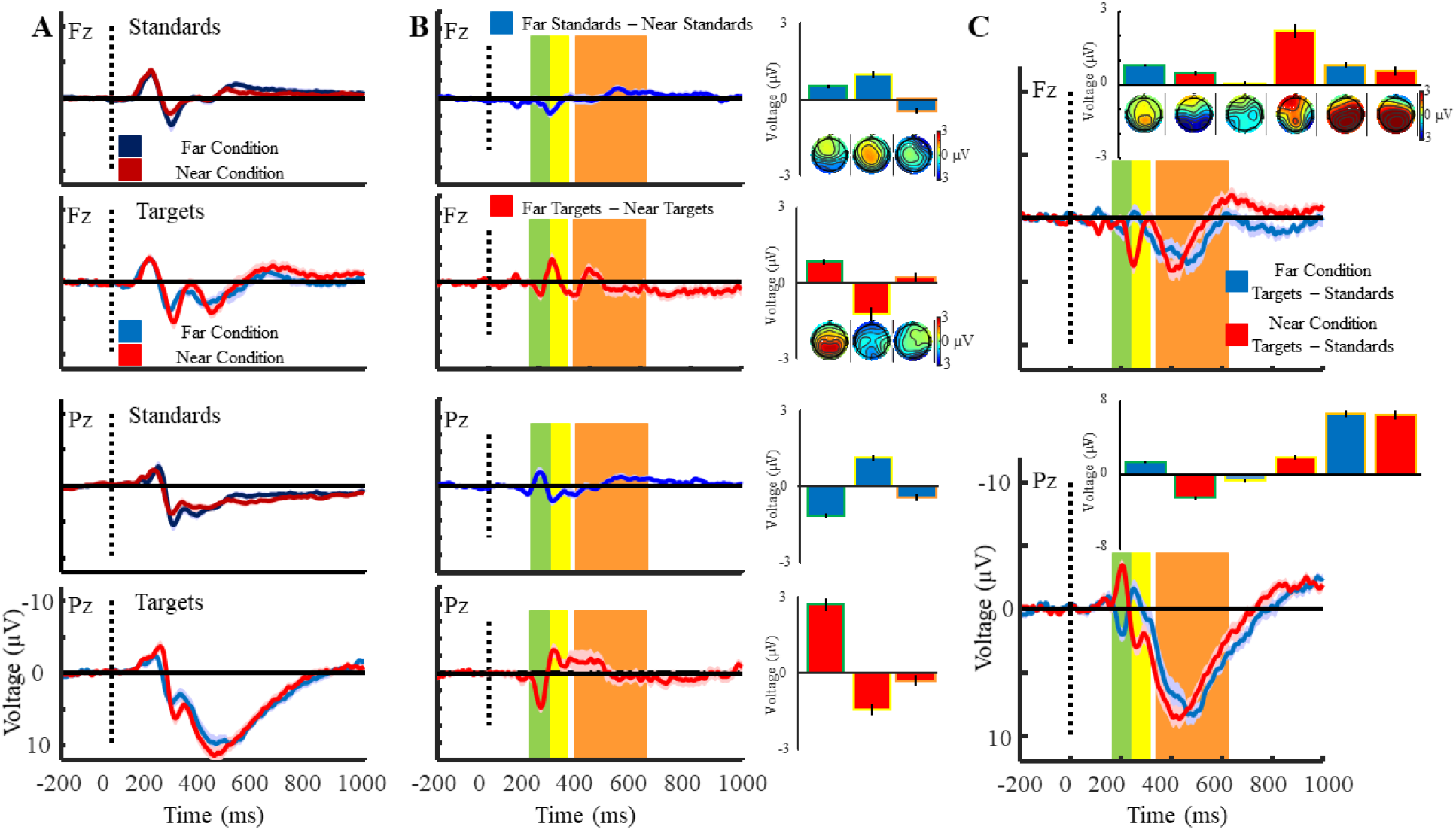
A) Grand-averaged ERP plots for target and standard stimuli at electrode Fz and Pz. B) Difference waveforms for near and far targets and standards. These waveforms were created by subtracting near standards from far standards, and near targets from far targets, at electrodes Fz and Pz. C) Difference waveforms for the Far Targets and Near Targets conditions. Plots were generated by subtracting ERPs to standards from the ERPs to targets in each condition. Far Targets Difference Wave = Far Condition Targets – Standards; Near Targets Difference Waveform = Near Condition Targets – Standards. Coloured shaded regions in each difference waveform represent the three time windows used for analyses. These windows are based on two opposing patterns of activity, and the P3 waveform, observed in the Far Targets and Near Targets difference waveforms at electrode Pz. Yellow = 175-225 ms; Orange = 225-300 ms; Red = 349-623 ms. Topographies represent activity across the scalp while bar graphs represent activity at either electrode Fz or Pz, averaged across these time windows. All error bars in each plot represent the standard error of the mean.

Table 4 shows the results of the one-tailed *t*-tests, effect size measurements, and Bayesian one-tailed *t*-tests for the difference waveforms. Table 4A shows the results of a one-tailed t-test comparing the standard and target difference waveforms from Figure 6B to zero. A right or left tailed t-test was performed based on the mean voltage of each time window. The results suggest a significant difference between Near and Far conditions at the two early time windows but not the later, P3 time window. A similar pattern can also be observed between Near and Far targets with significant differences observed during the 175-225 ms and 225-300 ms time windows. Table 4B shows results for the difference waveforms generated from the Near and Far conditions. A left or right tailed *t*-test was used based on the mean voltage of each time window. Hedge’s *g* was used to estimate effect size, which was calculated using version 1.5 of the Measures-of-Effect-Size toolbox for Matlab (Hentschke & Stüttgen, 2011). Results suggest that a significant P3 response is obtained during the 349-623 ms time window for both difference waveforms. There is also a significant deviation from 0μV during the 175-225 ms and 225-300 ms time windows for the Near difference waveform.

**Table 4:** Results of one-tailed *t*-tests, Bayesian one-tailed *t*-tests, and estimated measures of effect size for the ERP waveforms at three time windows. Hedge’s *g* is used to estimate effect size. Bayes factors were calculated by testing if the waveforms at each time window was less or greater than zero, based on the mean voltage. BF_10_ indicates support for H1 over H0.

| Condition | Electrode/<br>Time Window | Mean<br>Voltage<br>(μV) | Standard<br>Deviation<br>(μV) | $t$ (17) | $p$ | Confidence<br>Interval | Hedge's $g$ | $BF_{10}$ |
| --- | --- | --- | --- | --- | --- | --- | --- | --- |
| A | Pz/175-225 ms | -1.17 | 2.07 | -2.40 | 0.014 | -Inf:-0.32 | -0.54 | 2.28 |
| A | Pz/225-300 ms | 1.15 | 1.93 | 2.52 | 0.011 | 0.35:Inf | 0.57 | 2.77 |
| A | Pz/349-623 ms | -0.18 | 1.29 | -0.61 | 0.28 | -Inf:0.35 | -0.14 | 0.29 |
| B | Pz/175-225 ms | 2.70 | 2.52 | 4.54 | <0.01 | 1.67:Inf | 1.02 | 111.00 |
| B | Pz/225-300 ms | -1.44 | 2.91 | -2.10 | 0.25 | -Inf:-0.25 | -0.47 | 1.43 |
| B | Pz/349-623 ms | -0.53 | 3.25 | -0.70 | 0.25 | -Inf:0.80 | -0.16 | 0.30 |

A = Far Standards – Near Standards; B = Far Targets – Near Targets
| Condition | Electrode/<br>Time Window | Mean<br>Voltage<br>(μV) | Standard<br>Deviation<br>(μV) | $t$ (18) | $p$ | Confidence<br>Interval | Hedge's $g$ | $BF_{10}$ |
| --- | --- | --- | --- | --- | --- | --- | --- | --- |
| C | Pz/175-225 ms | 1.37 | 2.26 | 2.57 | <0.01 | 0.44:Inf | 0.58 | 3.02 |
| C | Pz/225-300 ms | -0.62 | 3.29 | -0.79 | 0.22 | -Inf:0.73 | -0.18 | 0.32 |
| C | Pz/349-623 ms | 5.77 | 3.11 | 7.88 | <0.01 | 4.50:Inf | 1.77 | 3.73e+4 |
| D | Pz/175-225 ms | -2.51 | 1.94 | -5.49 | <0.01 | -Inf:-1.71 | -1.24 | 641.08 |
| D | Pz/225-300 ms | 1.97 | 2.68 | 3.12 | <0.01 | 0.87:Inf | 0.70 | 7.87 |
| D | Pz/349-623 ms | 6.12 | 2.48 | 10.47 | <0.01 | 5.10:Inf | 2.36 | 1.52e+6 |

C = Far Condition Targets – Standards; D = Near Condition Targets – Standards

Results from the paired sample *t*-tests, estimates of effect size, and Bayesian paired sample *t*-tests are shown in Table 5, comparing the Near difference waveform to the Far difference waveform. Results suggest that our difference waveforms are not significantly different at the 349-623 ms time window but do differ during the 175-225 ms and 225-300 ms time windows. For the Far Targets – Near Standards difference waveform a positive deflection is observed during the 175-225 ms time window followed by a negative deflection during the 225-300 ms time window. An opposing pattern of activity is observed for the Near targets – Far standards difference waveform with a negative deflection during the 175-225 ms time window followed by a positive deflection during the 225-300 ms time window. Such patterns of activity can also be observed for the standard and target difference waveforms at electrode Pz, as shown in Figure 6B. Contrary to the ERP plots in Figure 6C, our conditions do not differ during the 225-300 ms time window. The above differences are also consistent with the results presented in Table 4.

**Table 5:** Results of two-tailed *t*-tests, Bayesian paired *t*-tests, and estimates of measure of effect size between the ERP waveforms at three time windows for electrode Pz. Hedge’s *g* is used to estimate effect size. Bayes factors were calculated by comparing both waveforms at each time window. BF_10_ indicates support for H1 over H0.

| Oddball Tasks Compared | Electrode/ Time Window | Mean Voltage (μV) | Standard Deviation (μV) | *t* (17) | *p* | Confidence Interval | Hedge’s *g* | BF<sub>10</sub> |
| --- | --- | --- | --- | --- | --- | --- | --- | --- |
| C:D | Pz/175-225 ms | 3.88 | 3.68 | 4.47 | <0.01 | 2.05:5.70 | 1.80 | 97.33 |
| C:D | Pz/225-300 ms | -2.59 | 4.03 | -2.72 | 0.014 | -4.59:-0.58 | -0.84 | 3.91 |
| C:D | Pz/349-623 ms | -0.35 | 3.44 | -0.43 | 0.67 | -2.06:1.36 | -0.12 | 0.26 |

**C = Far Condition Targets – Standards; D = Near Condition Targets – Standards**

Figure 7 shows a plot of the grand-average ERPs at the HEOG, which is designed to monitor horizontal eye movements. These plots were generated to observe changes in eye movements due to changes in the depths of the stimuli; diverging eye movements were expected for far stimuli and convergent eye movements were expected for near stimuli. Plots of the grand-average standards and targets difference waveforms (as shown in Figure 6) are also shown to compare HEOG changes to ERP changes. A two-tailed t-test was performed at each time point of the HEOG and Pz ERP to understand which time points significantly differ from 0, with significant tests indicated by pink marks at the bottom of each plot. For these t-tests, α was set to 0.005. For standards, there is no significant point of difference for the HEOG but for the Pz difference waveform the earliest point occurs 253 ms post-stimulus. For target stimuli, the earliest significant point of difference is at 179 ms for the HEOG and 190 ms for the Pz difference waveform.

**Figure 7:**
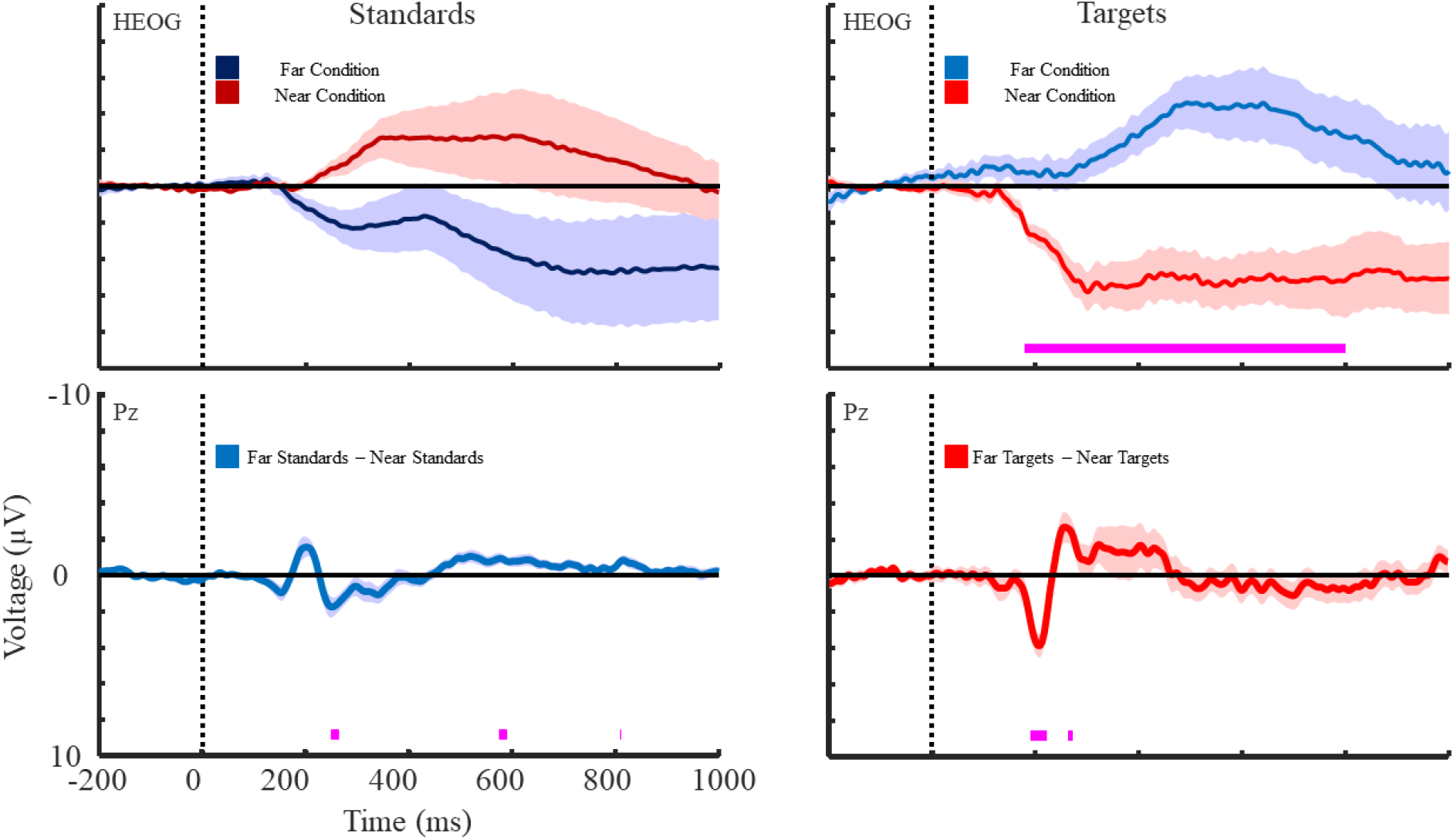
Grand-averaged waveforms following targets and standards in the Near and Far conditions for the HEOG and Pz electrodes. Pink regions along the x-axis represent areas where the Near and Far condition ERPs are significantly different. Error bars represent the standard error of the mean.

## 8. Discussion - Exp 2

To further demonstrate the ability of VR as an experimental tool, we had participants complete a novel, depth-based oddball task. The VR head mounted display provides several advantages compared to a non-VR monitor regarding EEG experimentation such as independent presentation of stimuli to each eye. During the depth oddball task, participants would complete two separate conditions, responding either to the near or far target and withholding responses to the respective standard. Separately presenting stimuli to each eye allowed us to manipulate the sense of depth while also maintaining the angle and subjective size of each orb from the participant’s point of view. Using this paradigm, we were able to demonstrate that early ERP components are sensitive to changes in stimulus depth while also eliciting the oddball-related P3 waveform.

Beyond a P3 response to targets at both depths, there were several interesting characteristics when comparing the Near and Far conditions. Comparing difference waves at Pz and Fz of the Near and Far conditions depicts an interaction of depth with elicited ERPs. Figure 6B summarizes these differences, particularly at Pz, where the early waveform components at 175-225ms (first window) and 225-300ms (second window) vary significantly for both Standard and Target difference waves. Standards show a negative deflection in the first window and a small positive deflection in the second, while targets show the opposite with a large positive deflection in the first window and a negative deflection in the second. These exciting results also show that ERPs differ with depths, supporting the findings of Liu, Meng, Wu and Dang (2015). However, our experiment likely doesn’t account for all interaction between the oddball stimulus and depth; further study with an extended experimental design is warranted. One such study may be a passive perceptual experiment involving all non-oddball elements at both depths, from this we would be able to gauge variance across ERPs that is only due to depth and not task demands.

Our current results support distinct physiological processes for interpreting stimuli across varying depths. The HEOG deflections portray an effect where the eye converges to focus on a near object and diverges to focus on a far object, which is expected. What was not previously expected was seeing the timing of this movement consistently appear alongside the ERP changes associated with that object’s onset. These novel results are interesting not simply due to the eye movement that is consistently observed with the onset of these objects, showing that depth cues are being processed by the eyes, but they also serve to show that there is a temporal alignment between the brain activity and eye movement. Further investigation of how that depth information processing produces cognitive and behavioural effects is needed.

However, the differential processing of two neutral stimuli that have been size-matched on the retina is not necessarily intuitive. Henderson, Vo, Chunharas, Sprague and Serences (2019) demonstrated that different depths are processed in different retinotopic areas of the brain, mainly in area V3A, and that an encoding model can be used to map the depth of objects on a z-axis, along with their positions in 2D space along vertical and horizontal axes. This provides an explanation of the different depths producing different ERPs. While this explanation points out a cognitive mapping perspective on where brain activity changes to these stimuli, it does not identify why such a mapping would be designed or beneficial. In that context, a possible explanation of this phenomenon is the evolutionary relevance of an object’s distance to the viewer; responding quickly to a stimulus that is near you tends to be more important than responding quickly to stimuli farther away. We may be physiologically poised to react faster to threatening stimuli that are closer, as seen in the earlier onset of eye movements in HEOGs for near stimuli when compared to far stimuli. The current study may present a generalization of this propensity in detecting and responding to non-threatening stimuli.

Another potential explanation is that differences between Near and Far difference waveforms may depict a perceptual crossing of a distinguishing boundary between peripersonal space (PPS) and extraperipersonal space (EPS), as described by di Pellegrino & Làdavas, 2015; Làdavas, 2002). PPS refers to a representation of space around an individual that is both malleable and reactive to task demands. There is evidence of distinct regions within this space (de Vignemont & Iannetti, 2015), and known factors demonstrated to change or modify the personal space (Canzoneri et al., 2013). Differences between the Near and Far conditions, as shown in the difference waves of Figure 6C, may be interpreted as an interaction with distinct regions of peripersonal space – near objects could be perceived to be within PPS and thus is more reactive than a far object that could be perceived to land in EPS.

A possible confound for the differences seen in the P3 of the Near and Far stimuli could be a difference in the complexity of the environment being perceived by the participants, as they could be processing the background and surrounding environment to a greater detail. While this is a valid concern, we don’t believe that the complexity of the environment or the background in the Far condition is in any way different to that of the Near condition, given that the participant’s position in the environment is identical in both conditions and the same environment is visible to them between all trials of all conditions. What we do propose as a possible explanation for these differences is that following the participant’s focus on the fixation cross, the onset of the Far stimulus would happen after a very brief time period where there is a retinal disparity of the far away object, which is then corrected by the eye to focus on the far object. This retinal disparity is mainly attributed to the difference in depth of the object and not the environment surrounding it, leading us to believe that once again the depth effect explains the differences seen in our P3 responses between the two conditions.

With all of these observations and proposed explanations in mind, we must also consider possible interferences and unintended behaviours that could confound these findings. Eye movements that are elicited from stimuli other than the objects presented could interact with the effects seen with the HEOGs. Changes in the exact viewing experience of each participant due to potential image blur, visual acuity, and headset positioning on head shapes and sizes that are inherently different between participants are other potential sources of variability. However, measures were taken to minimize these effects, such as adjusting the interpupillary distance of the lenses in the headset, verbal instructions to maintain eye position on the fixation cross, elimination of stimulus shadows and other such depths cues, and readjusting headset size to best fit each participant.

## 9. Conclusions

Our finding of similar ERPs for traditional and VR tasks in Exp. 1 show the effectiveness of using a VR head mounted display for ERP research. These results also demonstrate the generalizability of the P3 response, as well as its relevance and applicability in VR. We were also able to demonstrate the flexibility of VR in constructing a novel version of the oddball task, with near and far stimuli eliciting a P3 response along with depth-based differences earlier in the ERP waveform. This research helps to establish the utility of VR in creating new experimental paradigms within a controlled, laboratory setting.

If results can be validated with other tasks without participants being confined to a chin rest, assuming muscle artifacts can be properly accounted for, there may be a basis for conducting indoor mobile VR experiments. Leveraging recent advances in mobile EEG (Aspinall, Mavros, Coyne, & Roe, 2015; Zink, Hunyadi, Huffel, & Vos, 2016; Scanlon et al., 2019) with advances in the wireless VR industry, there may be space to harness the combination of VR with more mobile experiments within the laboratory.

To investigate the findings from Exp. 2 regarding differences in the Near and Far difference waves, we are currently running a similar experiment in which we present to the participant five size-matched orbs at different depths. The goal is to determine if there is a gradient-like depth effect, such that we expect the three additional depths to fall sequentially between the nearest and farthest orbs. In future studies, there is room to investigate how other domains interact with the depth oddball. Results may mirror comparable studies of overlapping visual and auditory modalities in the oddball task (Brown, Clarke, & Barry, 2007; Campanella et al., 2010; Robinson, Ahmar, Sloutsky, Robinson, & Sloutsky, 2010). Brown et al. (2007) showed how auditory and visual oddballs are paired to describe two distinct stages of auditory processing, an early modality-dependent stage and a later context-dependent stage. In addition to auditory dual tasks, we may integrate other visual domains in the depth oddball task such as colour, for example. This method would allow us to ascertain the flow and integration of depth specific information in concert with other modalities.

In line with research into peripersonal space, there is the opportunity to investigate depth-oddball sensitivity to known modulatory factors of peripersonal space, such as gauging the influence that target threat may have on the relative discrimination of near and far stimuli. In other words, how would different threats affect one’s mapping of peripersonal space? Depending on the threat content of the stimuli at various depths, there is reason to believe we could activate different PPS maps altogether (de Vignemont & Iannetti, 2015). One paradigm to introduce an element of threat while maintaining neutral stimulus presentation would be to offer a monetary sum at the beginning of the experiment. A loss is then accrued from the initial sum if response times fall below a certain baseline-derived threshold. A comparison of adherence to thresholds and resultant ERPs for each Near and Far conditions both with and without threat may reveal interactions of threat with different regions of personal space. The increased flexibility provided by VR systems has the potential to increase the breadth of EEG studies that can be conducted, greatly improving the external validity of EEG experimentation.

Here we show that a VR head mounted display is a great tool in developing a wide variety of preexisting experimental paradigms. Some topics that may readily benefit from a three-dimensional environment include visual attention, working memory, and learning, allowing for more accurate and realistic models of human navigation. VR as a tool in brain imaging studies offers increased generalizability and flexibility compared to traditional non-VR monitors This is achieved without sacrificing experimental control or introducing highly variable noise artifacts common with using EEG in environments outside of the laboratory.

## Acknowledgements

This work was supported by a discovery grant to KEM from the Natural Sciences and Engineering Research Council (NSERC) of Canada and start-up funds from the Faculty of Science. Thank you to all members and volunteers of the Mathewson lab for assisting with data collection and experimental setup. Thank you to our collaborators at IBM Research for providing help and guidance designing the virtual reality task and environment.

